# Postmortem MRI of Tissue Frozen at Autopsy

**DOI:** 10.1101/2024.01.20.576456

**Authors:** Govind Nair, Roy Sun, Hellmut Merkle, Kyra Hoskin, Kendyl Bree, Stephen Dodd, Alan Koretsky

## Abstract

**Introduction:** Postmortem MRI provides insight into location of pathology within tissue blocks, enabling efficient targeting of histopathological studies. While postmortem imaging of fixed tissue is gaining popularity, imaging tissue frozen at the time of extraction is significantly more challenging.

**Methods:** Tissue integrity was examined using RNA integrity number (RIN), in mouse brains placed between -20 °C and 20 °C for up to 24 hours, to determine the highest temperature that could potentially be used for imaging without tissue degeneration. Human tissue frozen at the time of autopsy was sealed in a tissue chamber filled with 2-methylbutane to prevent contamination of the MRI components. The tissue was cooled to a range of temperatures in a 9.4T MRI using a recirculating aqueous ethylene glycol solution. MRI was performed using a magnetization-prepared rapid gradient echo (MPRAGE) sequence with inversion time of 1400 ms to null the signal from 2-methylbutane bath, isotropic resolution between 0.3-0.4 mm, and scan time of about 4 hours was used to study the anatomical details of the tissue block.

**Results and Discussion:** A temperature of -7 °C was chosen for imaging as it was below the highest temperature that did not show significant RIN deterioration for over 12 hours, at the same time gave robust imaging signal and contrast between brain tissue types. Imaging performed on various human tissue blocks revealed good gray-white matter contrast and revealing subpial, subcortical, and deep white matter lesions typical of multiple sclerosis enabling further spatially targeted studies.

**Conclusion:** Here, we describe a new method to image cold tissue, while maintaining tissue integrity and biosafety during scanning. In addition to improving efficiency of downstream processes, imaging tissue at sub-zero temperatures may also improve our understanding of compartment specificity of MRI signal.

## Introduction

Human brain is often flash-frozen at the time of autopsy for long-term storage, which robustly preserves the tissue microstructure as well as DNA, RNA, and protein functionality.^1,2^ Ideally, brain is dissected into slabs of about 1-2 cm thickness and rapidly frozen by immersion in 2-methylbutane (an agent with high thermal conductivity), pre-cooled by placing over dry ice or liquid nitrogen, between temperatures of -70 °C and -160 °C.^3-6^ While snap-freezing the tissue reduces the possibility of freezing artifacts, and long-term storage is often done at -80 °C,^7^ the optimal temperature for sectioning brain tissue is much higher between -7 °C to -10 °C.^8^

Specific pathological regions can be targeted for immunohistology if their locations are known *a priori*. Efficient targeting of such regions could be achieved through imaging, either just before death, between death and autopsy (*in situ*), or of the postmortem tissue,^9^ and blocks can be precisely sectioned at the time of autopsy.^10^ However, specific circumstances often make imaging before death difficult, and *in situ* imaging can increase the postmortem interval (PMI), which can be detrimental for tissue preservation. Imaging fixed organs and tissue blocks has proved to be an invaluable tool for targeted histopathology in various neurological disorders and traumatic brain injuries.^10-14^ Similar imaging of frozen tissue blocks could decrease cost and time involved in spatially specific studies of molecular function, microstructural and immunocytochemical changes, and in situ hybridization studies. Imaging fixed human tissue has also been useful for understanding origins in-vivo MRI contrast in normal and disease states, such as myeloarchitecture and iron distribution.^15-20^

The main limitation to imaging frozen brains is the rapid decay of signal from protons in frozen tissue, making imaging difficult through traditional pulse-sequences.^21-23^ However, the freezing point of water within tissue depends on the compartment it resides in. Previous studies have shown signal from bovine tissue pieces extracted and frozen to -16.6 °C using short echo times of up to 1.1 ms in a 1.5T clinical scanner.^23,24^ Such studies demonstrate the feasibility of using non-solid state imaging techniques to image frozen tissue. However, routine imaging of cold tissue blocks would necessitate determination of optimal temperature for imaging, which produces high enough signal- and contrast-to-noise ratios (SNR and CNR) to complete imaging within a reasonable timeframe while maintaining tissue integrity and minimizing protein and nucleic acid degradation.^25^

Here, we describe a setup for MRI of tissue frozen at the time of autopsy while preserving RNA integrity, which serves as a robust and sensitive marker for tissue degeneration. As a first step, RNA degradation was measured from mouse brain tissue as a function of time at various temperatures. Then MRI setup, designed for imaging frozen tissue at the optimal temperature, was used to image the frozen tissue blocks. RNA integrity numbers (RIN) were determined before and after imaging of the cold tissue blocks to demonstrate the feasibility.

## Methods

To verify the optimal temperature range for conducting scans while preserving molecular-scale integrity, RIN analysis was performed on mouse brain tissue (n=8, age=12-month-old). Brains were dissected immediately upon euthanasia using 10% CO2 and excess liquid was blotted off the brain to minimize ice formation. Each hemisphere was added to a separate microcentrifuge tube, filled with isopentane and pre-cooled using dry ice. Any long-term storage of tissue was done at –80 °C. 25 mg blocks of tissue were placed in new tubes and kept at –20 °C, –5 °C, 0 °C, +4 °C, 20 °C for 1, 4, 6, 8, 12, 16, 20, and 24 hours. Following incubation, RNA samples were prepared using RNeasy kits (QIAGEN, Hilden, Germany). Briefly, 25-30 mg of mouse brains were isolated, placed in individual nuclease-free 2 mL centrifuge tubes (DNA LoBind® Tubes, Eppendorf) and homogenized using a rotor-stator homogenizer (TissueMiser, Fisher Scientific) for 60 seconds, resting between 20-second pulses. The RNA Integrity Number (RIN) of extracted total RNA was analyzed using Automated Electrophoresis System (TapeStation 4150, Agilent Technologies, Santa Clara, CA). Samples were spun down, 350 µL of the supernatant diluted and spin columns placed in collection tubes (RNeasy mini kit), spun down for 15 s at 8,000 g and eluted 3 times by adding 700 µL Buffer RW1 (Qiagen) and spinning. Samples were spun down at 8,000 RPM for one minute to elute RNA. To ensure samples were within the working range of the RIN assay, RNA concentration was verified using UV spectrophotometry (NanoDrop 2000, Thermo Scientific). 1 µL of eluted RNA and 5 µL RNA ScreenTape sample buffer was mixed in an 8-well PCR tube strip (Agilent Technologies) via vortexing for one minute (IKA MS3 Vortexer) and spun down for one minute (MyFuge C1012, Benchmark Scientific). Samples in 8-well tubes were heat-shocked for three minutes at 72 °C (C1000 Touch Thermal Cycler, Bio-Rad), placed on wet ice for two minutes, and spun down. Samples were placed in the Agilent 4150 Bioanalyzer for automatic quantification and RIN calculation using RNA ScreenTape (Agilent). RIN obtained were fit to monoexponentially decaying curve using Prism (GraphPad Inc, version 9.3) with plateau set to 0.

Frozen human brain slabs (n=3) were obtained postmortem from donors diagnosed with multiple sclerosis (originally processed at Rocky Mountain Tissue Bank, Aurora, CO and transfered to Daniel S Reich), and imaged one at a time. A frozen tissue block was placed inside a 3D-printed (Accura ClearVue resin printer) tissue chamber (Fig 1A) filled with 2-methylbutane (MilliporeSigma, Burlington, MA), and secured using thumb screws and silicone O-ring (with wide temperature range from -54 °C, 1/16 Fractional Width, 037, McMaster-Carr) for transportation and imaging. The tissue chamber was covered in absorbent pads and placed overnight in a portable freezer (LionCooler Pro Portable Fridge Freezer, With Battery, Acopower, Walnut, CA) set at -20 °C for equilibration. The schematic of the imaging setup is shown in Fig 2B. A recirculating chiller (MX07R-20-A11B Refrigerated Circulator, The Lab Depot, Dawsonville, Fig 1Bi in schematic) placed outside the magnet room circulates 50% v/v aqueous ethylene glycol mixture through the cooling chamber holding the tissue chamber (Fig 1Bii) and RF coil placed at the MRI (Fig 1Biii) isocenter. Insulated tubes (1/2” Insulated Chiller Translucent Silicone Tubing, Across International, Livingston, NJ) were used for circulating the refrigerant entering the MRI room through waveguide (Fig 1Biv). The cooling chamber (Fig 1C) was 3D-printed to match the tissue chamber dimensions, allowing thermal contact on 3 sides, and insulated on all other sides. Initial validation of temperature of 2-methylbutane in the tissue chamber in MRI was performed using fiberoptic temperature sensor (FOTEMP4 with FO Temp Sensor, TS5-20MM-06, measuring range = -200 °C to +200 °C, Micronor Sensors, Inc, Ventura, CA).

**Figure 1:**
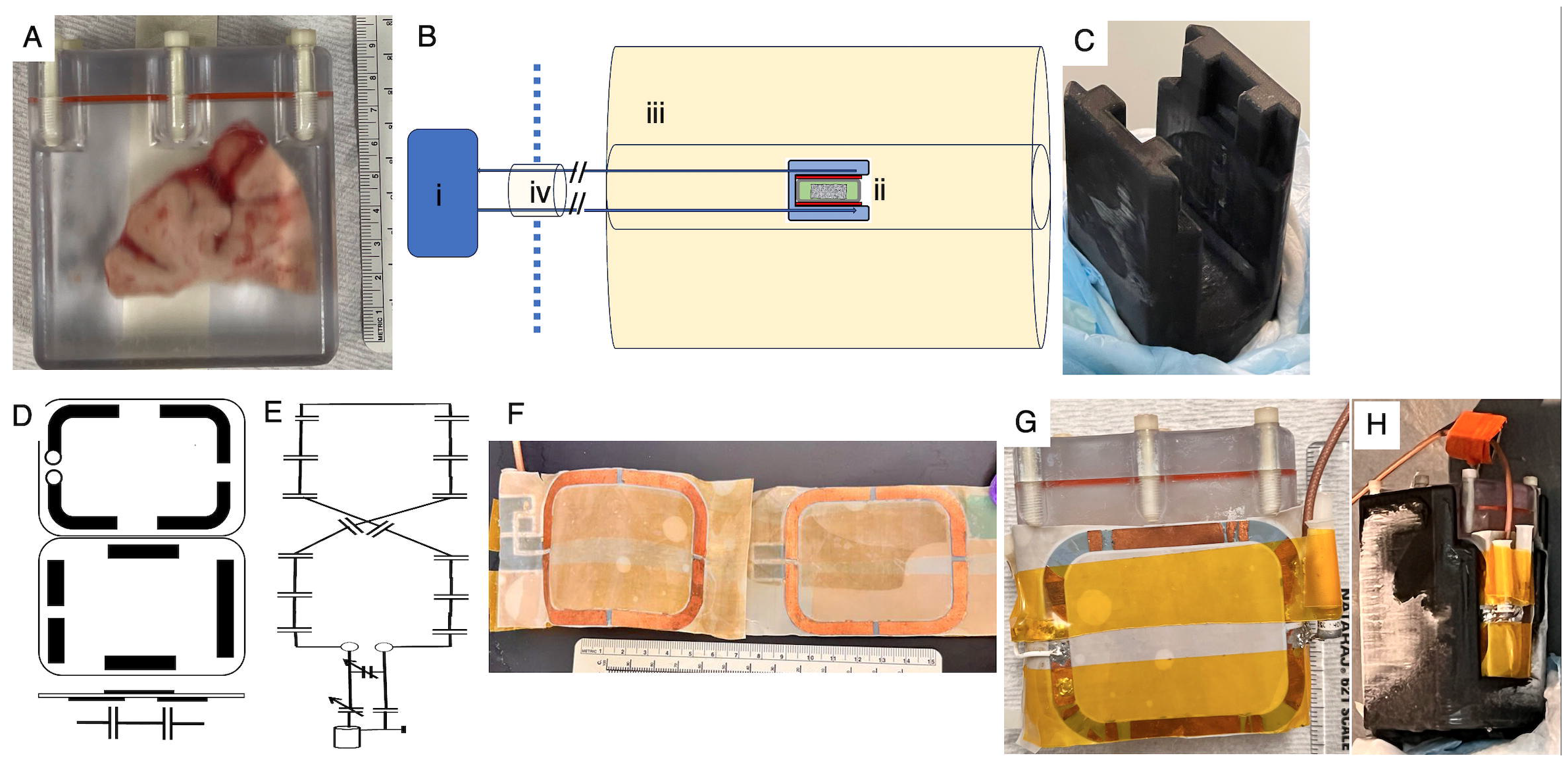
Details of MRI setup showing (A) brain tissue inside 3D-printed tissue chamber filled with 2-methylbutane and secured with thumb screws and O-ring (red); (B) schematic diagram of the setup to image cold brain tissue slabs with (i) recirculating chiller, (ii) cooling tissue chambers and RF coil, (iii) MRI, and (iv) waveguide with refrigerant tubes; (C) 3D-printed cooling chamber with insulating pads; (D) schematic diagram and (E) circuit diagram (F) photograph of the quasi-Helmholtz RF coil; (G) and photographs the placement of the RF coil around the tissue chamber and (H) which is them placed inside the cooling chamber.

**Figure 2:**
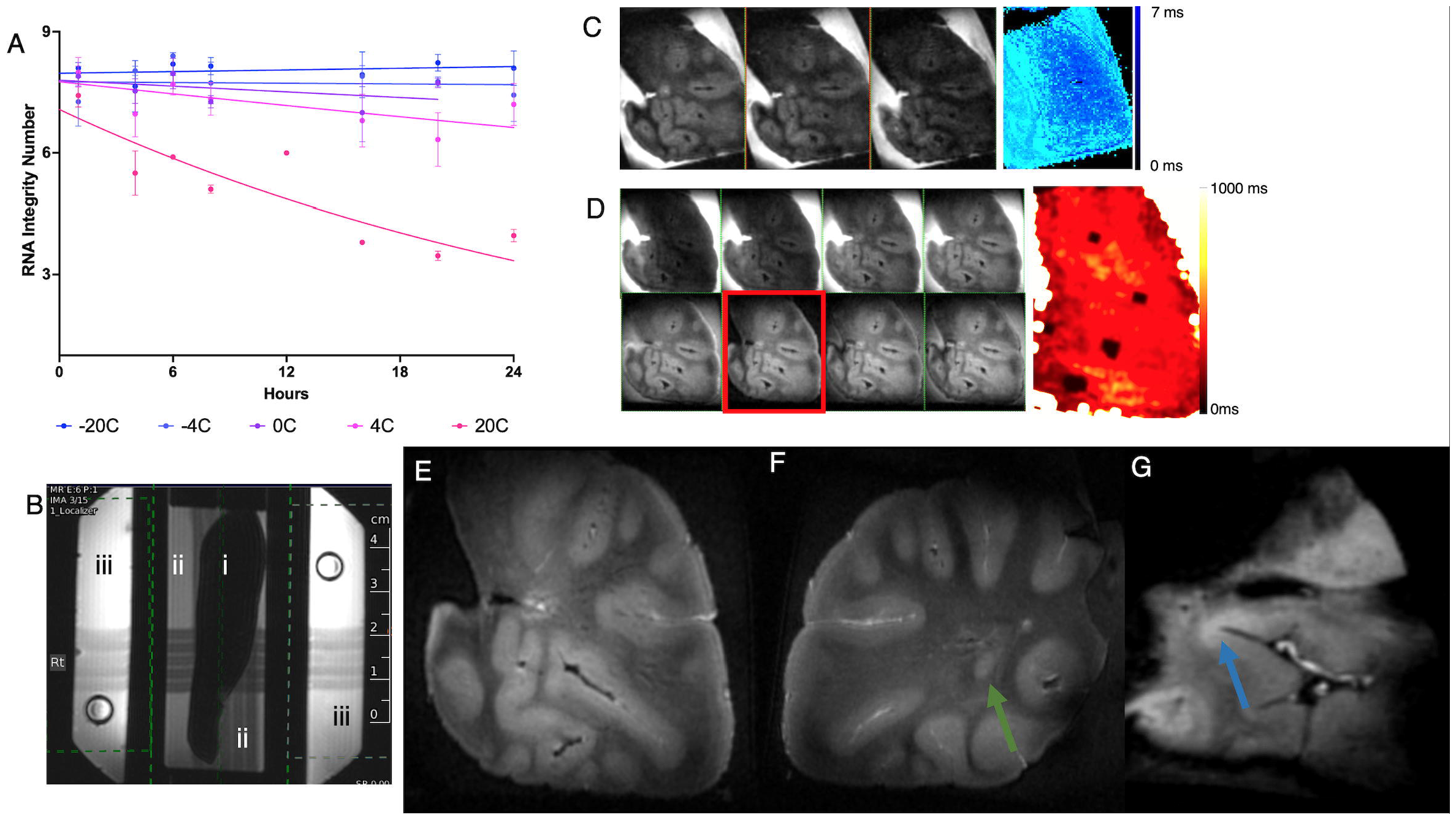
(A) RNA Integrity Number (RIN) determined from tissue placed at various temperatures (different colors) for various amounts of time, and best monoexponential decay fit of the RIN with time for various temperatures (various lines). (B) MRI of the setup depicting (i) tissue, (ii) tissue chamber, and (iii) cooling chambers on a localizer image. (C) Multi-echo GRE acquired at various TE (gray scale images from left to right 0.9 ms, 1.5 ms, 2.4 ms), and T_2_-map (in blue-light blue color scale) of a cold human brain slab. (D) MPRAGE images acquired at various Inversion Times (gray scale, top row left to right 50 ms, 200 ms, 400 ms, 600 ms and bottom row left to right 1000 ms, 1400 ms, 1800 ms, 2000 ms TI), with T_1_-map shown in color. 2-methylbutane signal is completely nulled at TI=1400 ms (red box in D). (E-G) MPRAGE images acquired at 375 µm isotropic resolution from three different brain slabs showing gray-white contrast, white matter lesion (green arrow), and suspected cortical lesion (blue arrow).

Imaging was performed on a 9.4T Bruker Biospec MRI system (Bruker BioSpin) in the MIF with custom build thin quasi-Helmholtz RF coil, schematic of which is shown in Fig 1D, and circuit diagram in Fig 1E. The transmit/receive coil was carved from double layer copper plated Teflon (CuFlon^®^ Microwave Substrates, Crane Co., West Caldwell, NJ) with a thickness of ∼0.26 mm, with ∼0.1 mm thickness copper (3 oz/square inch) on each side to fit between the tissue and cooling chambers. Two 50 × 64 mm (inside measurement) rectangular loops with rounded corners were carved, and segmented, using multiple gaps in a way that upper- and lower-side tracks overlapped partially forming multiple in-line capacitors along the loop (Fig 1F). The loops were connected in series and placed on top and bottom of the tissue container, therefore defining a figure-8 quasi-Helmholtz structure. Loops were terminated to the transmission line with a regular tune/match circuit with variable capacitors (Voltronics NMNAJ). The RF coil is housed between a cooling and tissue chambers (Fig 1G, H), thereby protecting it from condensation at the low imaging temperatures.

T_1_-mapping was done using 3D MPRAGE sequence with a TR=4000 ms, TE=1 ms, TI varied between 50 and 2000 ms in uneven steps, image resolution of 375 µm, and acquisition time of 11 minutes per TI. Similarly, 3D GRE sequence was acquired with a TR= 40 ms, TE=0.9-2.4 ms for T_2_-mapping. Voxelwise three-parameter fitting was performed in MATLAB (R2022a, Mathworks, Natick MA) to yield the T1 and T2 maps. High resolution imaging was performed using MPRAGE sequence with TR=4000 ms, TE=1 ms, TI=1400 ms, 12 averages, isotropic resolution of 300-400 µm and a scan time of 4 hours. Tissue was typically in the MRI for a total duration of 6 hours, however longer total times of up to 12 hours were used during T_1_- and T_2_-mapping as well as SNR testing.

## Results

RINs were measured at various times for each temperature setting was fitted to a monoexponentially decaying function and the rate of decay determined (Fig 2A). No significant RIN deterioration was seen at -20 °C (k=-8.5×10^−4^ /h, 99% CI: [-5.2×10^−3^ /h, 3.5×10^−3^ /h], p>0.05), -4 °C (k=-3.7×10^−4^ /h, 99% CI: [-4.6×10^−3^ /h, 5.3×10^−3^ /h], p>0.05), or 0 °C (k=3.1×10^−3^ /h, 99% CI: [-2.5×10^−3^ /h, 8.9×10^−3^ /h], p>0.05) over 24 h. The experiment demonstrated an exponential decay in RIN at 4 °C (k=6.6×10^−3^ /h, 99% CI: [-6.1×10^−4^ /h, 1.2×10^−2^ /h], p<0.05) and 20 °C (k=3.1×10^−2^ /h, 99% CI: [2×10^−2^ /h, 4.3×10^−2^ /h], p<0.05), a rate which was significantly different from each other (p<0.05) (Fig 2A). The decay constant at 4 °C and 20 °C was significantly higher than that at -4 °C in the first 24 hours after tissue extraction (p<0.05). Given the RNA degradation rate and values at 12 hours, a temperature below -4 °C was chosen for imaging.

Tissue and cooling chambers were 3D printed, and a thin RF coil designed to fit between the tissue and cooling chambers. The coil tuning was found to be robust to changes in temperature between -10° C to 0 °C. There was a gain in temperature of about 5 °C in the refrigerant ethelyne glycol mixture as it flowed from the chiller located outside the magnet room to the MRI isocenter in the setup used. Preliminary tests revealed sufficient SNR at temperatures up to -10 °C (data not shown). Therefore, the recirculating chiller was set to -12 °C to maintain a safe operating temperature of an -7 °C within the tissue chamber for imaging. The signal from tissue (Fig 2B-i) at that temperature was about one-fiftieth of that at the temperature 2-methylbutane in the tissue chamber (Fig 2B-ii), necessitating suppression of the signal from the bath. Signal from ethylene glycol in the cooling chamber (Fig 2B-iii) was brighter than that from 2-methylbutane be a factor of 2, and frequency shifted by 3.5 ppm, which enabled the use of frequency-selective suppression.

MGE images acquired with variable TE from 0.9 ms to 2.4 ms revealed a T2 of 0.65 ms (95% CI of 0.45-0.84 ms, Fig 2C, b/w images showing images acquired at various TE, T2 map shown in color) in brain, while that in 2-methylbutane was not calculated. Separately, MPRAGE images acquired while varying TI from 50 to 2000 ms (Fig 2D, b/w images showing images acquired at various TI, T_1_ map shown in color) revealed an average T1 of 251 ± 52 ms in the brain and 2052 ± 100 ms for 2-methylbutane. The null point in signal from 2-methylbutane was 1400 ms, and that was used for all high-resolution images of the tissue. The inversion efficiency achieved in the average cold tissue was 0.45 while that in the 2-methylbutane was 0.98, indicating the challenges to accurate calibration of the RF pulse as well as T_2_ relaxation during the hyperbolic secant inversion pulse (8 ms duration used in these experiments).

High resolution MRI of cold brain tissue samples of patients who died with a clinical diagnosis of multiple sclerosis obtained from Rocky Mountain Brain Bank at 375 µm isotropic revealed anatomical details such as GM and WM boundaries, and vessels (likely due to residual blood or voids). In addition, white matter lesion (Fig 2 E and F, white arrows), and cortical lesions (Fig 2G, green arrow) could be identified.

## Discussion

Here, we demonstrate MRI of cold tissue maintained at temperatures that preserve RNA within the magnet. A recirculating chiller pumped refrigerant from outside the magnet room, into cooling chamber located at the magnet isocenter encasing biosafe tissue chamber, and MRI was acquired using novel, thin, quasi-Helmholtz RF coil placed between the two chambers. The tissue was maintained at temperatures that did not show RNA degradation within durations deemed necessary for imaging. Anatomical and pathological details could be clearly identified in the cold tissue with total time for tissue within the MRI ranging from 6-12 hours.

Use of tissue chamber sealed in a biosafety cabinet eliminated aerosol productions during the free handling of unfixed human tissue, which is essential for the human pathologies to be investigated using this setup. A portable freezer was used for overnight equilibration of tissue at intermediate temperature of -20 °C, as there is little evidence of RNA degradation at that temperature, and it enabled shorter times for equilibrating to the imaging temperature within the MRI scanner. The portable freezer also facilitated transportation from labs to MRI suite under sterile conditions. The recirculating cooling system was set up outside the magnet room, with refrigerant running into the magnet room in insulated line through the radiofrequency shield waveguides. The tube carrying refrigerant was never in contact with tissue or fluid that bathes the tissue, and imaging cradles and chamber could be cleaned with disinfectant or bleach.

Previously, imaging of cold tissue slabs has been attempted with thermal gradient achieved through dry ice or liquid nitrogen.^23,24^ Such a setup does not allow for precise control of tissue temperature and does not maintain sterile conditions within the magnet. A major limitation of the setup is that we did not directly monitor the temperature in the tissue chamber during scanning, which would have involved compromising the integrity of the tissue chamber to insert a thermocouple of optic fiber probe. Temperature monitoring could also be performed using infra-red sensors, frequency or phase shits of tissue or surrounding bath (MR thermometry), or through placement of silica gels directly within the tissue chamber, all of which would maintain the integrity of the tissue chamber.^21,26^ However, infrared remote temperature monitoring may require exposing a part of the tissue chamber, which could lead to further loss of cooling efficiency. Nevertheless, this is a feasible option and needs to be explored further. Absolute temperatures are not measurable using MR thermometry, and calibration involved in the processes could significantly increase the total scan time. Finally, temperature monitoring using silicone materials^21^ within the tissue chamber is an attractive option would need to be evaluated before adaptation.

Nevertheless, the temperature of the tissue chamber was monitored under the same conditions as a tissue scan but without human tissue in the tissue chamber using fiber optic probes. A consistent temperature differential of +5 °C was observed between the set temperature of the recirculating chiller and the 2-methylbutane in tissue chamber placed in the isocenter of the 9.4T MRI. This experiment would need to be performed as a quality assurance measure of the insulating efficacy of the various tubes and chambers. Indeed, it is important to allow the tissue to equilibrate to the temperature of the tissue chamber before MRI, which added about one hour to the total time the tissue was in the MRI.

We hypothesized that there may be sufficient pool of unfrozen water in tissue at sub-zero temperatures to provide tissue contrast with conventional imaging sequences. The proton NMR signal intensity has been shown to drop precipitously at 0 °C, and continues to drop exponentially with sub-zero temperature.^22-24^ There may be several factors leading to this observations, such as freezing of water in a tissue compartment (say extracellular vs. intracellular compartment), water existing in pockets of various sizes and pores within tissue, and a relatively low water exchange between pools as has been shown in various biological systems.^27-29^ The differences in T_2_ relaxation times between white and gray matter was sufficient to visualize anatomical structures and pathological regions in the MS brain slabs studied herein. However, it should be noted that method used for freezing the sample will result in considerable variation in the hydration, solute concentration, and cellular integrity,^30^ which may cause contrast variability between samples. Nevertheless, such compartment-selective signal may be used to better understand the in-vivo signal and improve its modeling for developing biologically relevant imaging markers.

Given the RIN degradation experiment and temperature gain expected in the setup, we chose to use a temperature of -7 °C for imaging, which was below the temperature to preserve RNA. Indeed, the long imaging times expected for high-resolution imaging of the tissue even with conventional pulse sequences necessitated this safety margin. Moreover, the T_1_ and T_2_ relaxation times of biological samples drop at high fields and in the temperature-range used here, causing significant challenges to using conventional imaging sequences. The short T_1_ relaxation times would yield little contrast between tissue types, and the short T_2_ relaxation times would cause a drop in signal at normally achievable TEs. Ultra-short or zero echo time (UTE or ZTE) sequence, while achieving TE in the micro-second range, are plagued with artifacts such as distortions and due to eddy currents. Image distortions caused by eddy currents would need to be corrected using post-processing before efficient histopathological targeting.^31^ Solid-state imaging, on the other hand, opens up more contrast opportunities by utilizing the chemical shielding and dipolar coupling, but can take very long to perform.^32^ Indeed, such sequences may also make visible some of the plastics used in creating the setup. Nevertheless, this technique can be implemented at even lower temperatures where the fraction of unfrozen water is lower or absent.

Maintaining cooler temperatures inside the magnet bore might be achieved by circulating super-cooled nitrogen gas into the cooling chamber if condensation within the magnet bore can be avoided. Since nitrogen gas would be MRI invisible, it allows more flexibility in imaging; MRI signal from the refrigerant used here, ethylene glycol, was suppressed using frequency and spatial saturation pulse. Indeed, immobilization of protons at lower temperatures cause an increase in T_1_, which might be useful in generating contrast between tissue types. Other considerations at lower temperatures include ice formation on the cooling and tissue chambers, signals from which would need to be suppressed, and versatility in RF coil for good performance at much lower range of temperatures.

A novel thin, pseudo-Helmholtz coil was constructed for the imaging performed. This construction method was chosen since space was very limited between sample and the cold finger structure that needed to be in tight thermal contact with the sample. In-line capacitors enabled building a relatively large coil at 9.4 Tesla (400 MHz). This approach also enabled stable operation using only minor adjustments despite some sample load variations, with reference power in the 2W range.

Indeed, images acquired on several blocks of postmortem tissue from patience once diagnosed with multiple sclerosis clearly showed grey, white matter, and lesions. The regions can easily and efficiently be targeted for further analysis since the shape of the tissue blocks do not change during imaging.

In conclusion, a method has been developed that enables MRI on previously frozen tissue to identify lesions within a tissue block without causing significant degradation of tissue. The setup is relatively straightforward and can be used on any MRI. This opens the possibility for prescreening the large number of human frozen tissues prior to histology to better guide histology or RNA analysis. In addition, the ability to vary temperature or to get tissue contrast at high resolution should open increased flexibility to understand MRI contrast relevant to normal anatomy and pathological states of the human brain.

## Acknowledgements

Study was funded by the intramural research program at the NINDS. We thank Daniel S. Reich and Martina Absinta for providing frozen human tissue blocks for developing this imaging system. We thank George Dold and Katherine Cameron, Section on Instrumentation, for assistance with design of tissue and cooling chambers.

## Author Contributions

Authors GN, SD, and AK contributed to contributed to conception and design of the study. Authors GN, RS, HM, KH, KB, and SD contributed to acquisition and analysis of data. Authors GN, RS, HM, AK contributed to drafting the text and preparing the figures. The authors declare no competing interests on the subject of the study.

## Notes

### Competing Interest Statement

The authors have declared no competing interest.

